# New clues about ribosome hibernation in microsporidia revealed by cryo-electron tomography

**DOI:** 10.1101/2025.08.26.670498

**Authors:** Mathew McLaren, Rebecca Conners, Michail N. Isupov, Vicki A. M. Gold, Bryony A. P. Williams, Bertram Daum

**Affiliations:** Living Systems Institute, University of Exeter, UK; Department of Biosciences, University of Exeter, UK; Biocatalysis Building, Department of Biosciences, University of Exeter, UK

## Abstract

Microsporidia are single-celled parasitic eukaryotes that alternate between a dormant environmental phase and a metabolically active, progeny-producing phase inside their animal hosts. A successful transition between these phases requires a tight regulation of translation, which must be suppressed during dormancy and rapidly reinitiated upon host ingress. Previous studies by us and others have shown that translational silencing in microsporidia is underpinned by ribosome hibernation, a process involving both the binding of hibernation factors and the formation of ribosome dimers. Using focussed ion beam milling and cryo-electron tomography to image dormant microsporidian spores, we have gained new structural insights into this hibernation mechanism. Our data reveal that ribosome dimers within the spores assemble into ordered arrays surrounding the convex surface of polar tube, a specialised organelle essential for host cell invasion. Sub-tomogram averaging resolved the structure of the spore-borne ribosomes, further extending our understanding of ribosome hibernation in microsporidia.

## Introduction

Microsporidia are obligate, intracellular eukaryotic parasites that exist as dormant spores in the environment ^1,2^. These parasites infect a wide range of animal species, including humans, and are transmitted primarily through the ingestion of contaminated water or food ^1,3^. In immunocompromised individuals, microsporidia cause a debilitating disease called microsporidiosis. In agricultural contexts, particularly in aquaculture across South-East Asia, microsporidian infections can devastate farmed livestock, leading to significant economic losses ^1,4^.

During their evolution, microsporidia have adapted to a specialised parasitic lifestyle. They are among the smallest eukaryotic cells, measuring only 2–5 µm in diameter, and have undergone extreme genome reduction, with genome sizes down to 2,300 kbp. As a result of this, Microsporidia also exhibit a minimalistic cell structure, lacking mitochondria and peroxisomes entirely, rendering them incapable of reproducing outside a host cell ^5–8^.

To infect host cells, microsporidia employ a specialised invasion organelle called the polar tube. Within the host’s gut, dormant spores germinate, explosively firing the polar tube in less than a second ^9,10^. This hollow tube, approximately 200 nm in diameter and extending dozens of micrometers in length, harpoons the host cell membrane and injects the infectious sporoplasm into the cell ^3,9,10^. Once inside, the parasite (now called meront) exploits the host’s metabolic resources to divide and differentiate into sporonts and finally, new spores ^2^. These spores eventually fill a large portion of the host cell and egress via lytic or non-lytic pathways. Some spores disseminate within the host’s body, while others are excreted to propagate the infection further ^2^.

During their dormant state in the environment, microsporidia must conserve energy to survive extended periods of time without a host. During this period, protein biosynthesis, one of the most energy-intensive cellular processes, must be minimised ^11^. In previous research, we utilised electron cryo-tomography (cryoET) of the microsporidian model *S. lophii* to investigate hibernating ribosomes in transit through germinated polar tubes ^12^. Studying these ribosomes via sub-tomogram averaging, we discovered that they form dimers. This was surprising, as hibernating ribosome dimers had until then only been seen in bacteria ^13–21^. However, we found that unlike bacterial hibernating ribosomal dimers (bHRDs), which use hibernation factors like Hpf and Rmf, microsporidian hibernating ribosome dimers (mHRDs) assemble via a distinct dimer interface that includes the ribosomal proteins eS31 and eS12 ^12^.

Within the mHRD of *S. lophii*, each ribosome is inhibited by the hibernation factor MDF1. This factor occupies the E-site of the ribosome, blocking the binding of tRNAs to the corresponding site, and drawing the L1 loop into a closed conformation. This renders the ribosome incapable of performing protein translation. This confirmed preceding cryoEM studies, where isolated hibernating monosomes from the microsporidia *Encephalitozoon cuniculi* and *Varimorpha necatrix* were also found to be inhibited by MDF1 ^11,22^. In *V. necatrix*, however, the polypeptide exit tunnel (PET) is additionally blocked by a second hibernation factor, MDF2 ^11^. Moreover, the hibernation factor Lso2 was identified in hibernating monosomes from *Paranosema locustae* ^23^, as well as in yeast ribosomes, where it simultaneously obstructs both the mRNA binding pocket, and the polypeptide exit tunnel ^24^.

The observed variability of hibernation factors extends into mammals, where multiple types of hibernating ribosomes have been identified. Ribosomes isolated from human cell cultures exhibit two distinct silencing states. One involves a non-rotated/non-ratcheted conformation bound by CCDC124 (the mammalian homolog of Lso2), along with EBP1 blocking the polypeptide exit site ^24^. The second state is rotated, where CCDC124 is replaced by SERBP1 and eEF2 ^24^. While CCDC124/SERBP1 and eEF2 function similarly by blocking the mRNA entry channel and inhibiting the A and P sites, EBP1 prevents interactions between stalled ribosomes and nascent polypeptide-associated factors, including RAC, SRP, Sec61, and NatA ^24^.

Another mammalian ribosome hibernation mechanism has been identified in reticulocytes, where 80S ribosomes are inactivated by IFRD2 (Interferon-Related Developmental Regulator 2) ^25^. IFRD2 binds to the P and E sites, using its C-terminal α-helix to sequester mRNA. Additionally, a deacylated tRNA occupies an alternative site beyond the E-site, termed the Z site ^25^. It has been proposed that this stably bound Z-site tRNA serves as a marker of stalled ribosomes, particularly in conditions of amino acid depletion or limited availability of translation factors ^25^. Biochemical studies and negative stain electron microscopy indicated that ribosomal dimerisation may also occur in rat cells during amino acid starvation ^26^. A high-resolution structure of a mammalian or any other eukaryotic hibernating ribosome dimer has yet to be determined.

Here, we employed Focused Ion Beam Scanning Electron Microscopy (FIB-SEM) combined with CryoET to investigate the structure and organisation of ribosomes within ungerminated *S. lophii* spores. We observed that mHRDs are positioned around the convex surface of the polar tube, with some tethered to its surface. Many of these tethered ribosomes are arranged in two-dimensional (2D) crystalline arrays. Using sub-tomogram averaging, we resolved the structure of the spore-borne mHRD at 6.5 Å resolution, identifying additional components involved in the dimer interface, as well as the ribosomal subunits that appear to maintain contact sites within the crystal.

Our findings expand our knowledge about the mechanisms of ribosome hibernation in eukaryotes in general and microsporidia in particular. Moreover, we propose a model wherein polar tube firing, which involves a topological eversion, triggers a sequential unpacking of the ribosomes, from crystals to dimers, likely culminating in active 70S monosomes as the parasite enters the host cell. This model provides new insights into how microsporidia optimise their ribosomal organisation and activity and how they rapidly activate ribosomes during host cell invasion.

## Results

To visualise hibernating ribosomes within dormant, ungerminated *S. lophii* spores, we isolated the species from monkfish (*Lophius piscatorius*) caught in the North Atlantic and landed in Brixham, UK. The spores were subsequently prepared for cryo-ET using two distinct methods: direct plunge-freezing on cryoEM grids or High-Pressure Freezing (HPF) using the Waffle method ^27,28^.

Following sample preparation, grids were transferred into an Aquilos cryoFIB/SEM (Thermo Fisher Scientific), where multiple lamellae were generated using a gallium-focused ion beam (FIB) (Fig. 1a,b). These milled grids were then transferred to a Titan Krios Transmission Electron Microscope (TEM; Thermo Fisher Scientific) for tomographic data acquisition. The resulting tomograms were reconstructed, segmented, and analysed, revealing the presence of dozens to hundreds of mHRDs within the majority of the tomograms. While many ribosomes were randomly distributed within the sporoplasm, a significant proportion lined the outer (convex) surface of the polar tube, which remained coiled within the spore (Fig. 1c). These ribosomes were organised into curved, 2-dimensional arrays. 3D classification (Supplementary Figure 2) revealed that while 70% of all ribosomes formed mHRDs, 30% were monosomes (Fig. 1d). However, the ribosome crystals appeared to solely consist of mHRDs.

**Figure 1.**
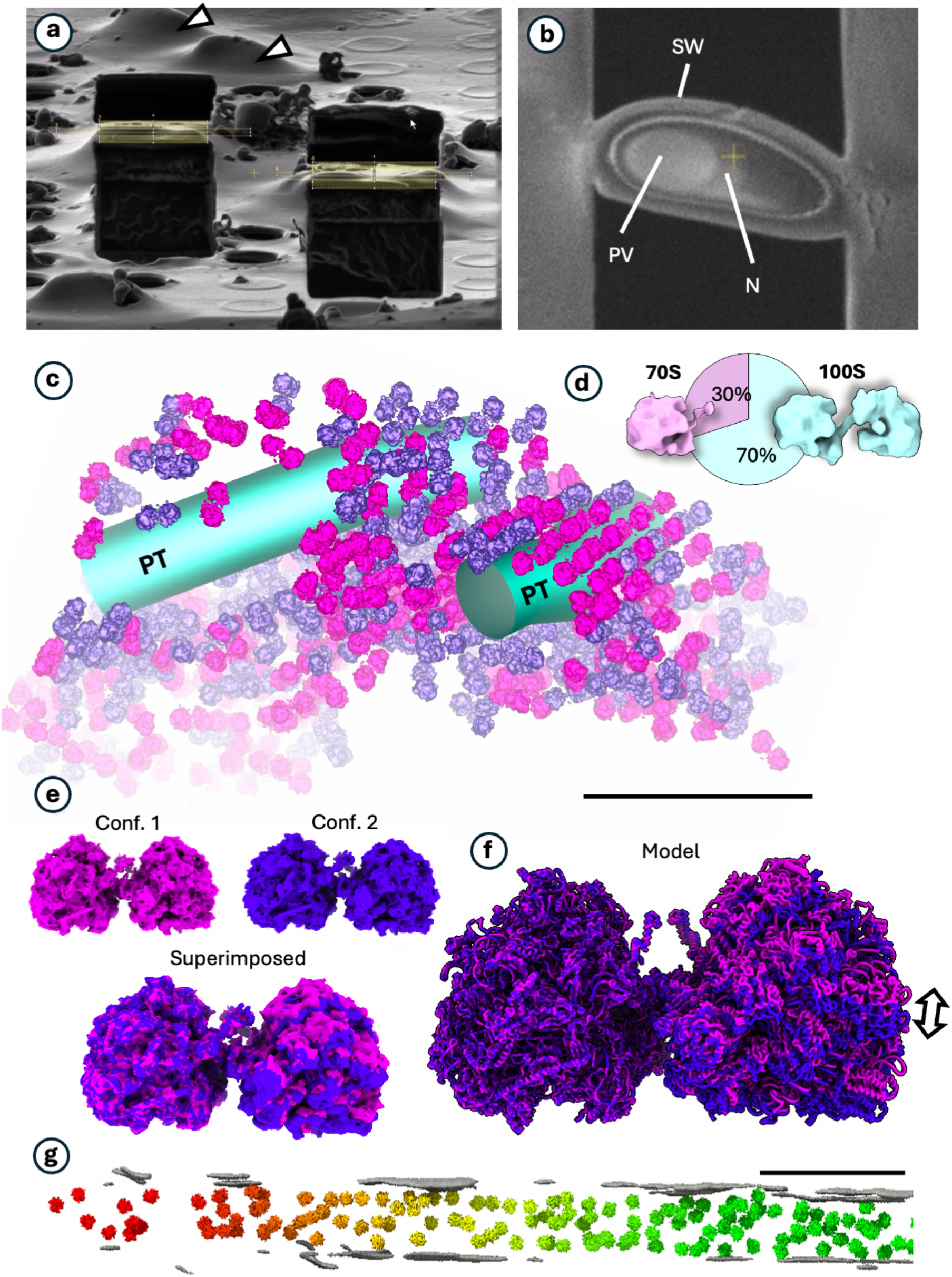
*In situ* organisation of mHRDs in *S. lophii* spores vs. fired polar tubes. **a**, tilted SEM view of two lamellae prior to thinning. White arrowheads indicate un-milled spores. **b**, SEM top view of a lamella through a spore. Spore wall (SW), posterior vacuole (PV) and nucleus (N) are indicated. **c**, segmented tomogram showing the polar tube (PT; teal) and mHRDs (purple and magenta). The two colours indicate two distinct classes of mHRDs. **d**, 3D classification reveals that 70% of all spore-borne ribosomes are dimers and 30% are monosomes. **e**, 3D classification further reveals two ribosome dimer conformations. These are shown separately (top panel) and superimposed (bottom panel). **f**, the two conformations shown as superimposed atomic models (liquorish representation). **g**, distribution of ribosomes within fired polar tubes of *S. lophii*. The ribosomes have relocated from the PT’s convex exterior into its interior lumen. The crystals have dissolved. Polar tube wall and membrane, grey; ribosomes, multicolour. Scale bars, 200 nm.

Moreover, 3D classification of the mHRDs in spores also revealed two distinct conformational classes (Fig. 1e, Movie 1, Supplementary Figure 3). These classes represented dimers with variations in the degree of bending at the dimer interface, specifically along an axis orthogonal to the two-fold symmetry (Fig. 1f). This shows that the dimer interface exhibits significant flexibility, explaining the difficulty of achieving high-resolution reconstructions of the entire mHRD complex. We then investigated whether differences in dimer angles relate to the curvature of the polar tube, which is contacted by the mHRDs and may dictate whether HRDs form crystalline arrays or remain as free-floating particles. To this end, we mapped the two distinct HRD conformations back onto their original tomograms. However, no clear correlation was observed between dimer angle and spatial distribution (Fig. 1c). Both conformational states were present in crystalline arrays as well as in the cytoplasm as free-floating dimers. This suggests that the dimer angle does not play a determining role in polar tube association or crystal assembly. The biological significance of this structural variability remains unclear and thus warrants further investigation.

CryoET of polar tubes after firing showed that ribosomes are now located inside the polar tube’s interior (lumen), in accordance with previous observations made by us and other groups (Fig. 1g, Supplementary Figure 4) ^12 29^. We previously determined the ratio of hibernating ribosome dimers to monomers in fired polar tubes is 70:30^12^ (Supplementary Figure 4), which exactly reflects the ratio in ungerminated spores (Fig. 1d). However, we do not find any vestiges of ribosomal arrays after polar tube firing. This is in line with observations made by Sharma *et al*, where ribosome arrays were observed in fired polar tubes, but in a minority of cases ^29^.

We next asked whether the structure of the spore-borne mHRDs differs from that of the ribosomes transiting through the polar tube following discharge ^12^. We therefore picked mHRDs from tomograms of spores and subjected them to sub-tomogram averaging using the Warp-Relion-M pipeline ^30^. This analysis yielded two conformational states of the dimer at 9.8 Å and 10.9 Å (Fig. 1e,f; Supplementary Figure 3). To improve the resolution of the mHRD structure, we sought to minimise the impact of the flexible dimer link. Thus, the 70S monosomes within each dimer were independently centred, masked, and averaged, yielding a 6.5 Å map for the half-dimer (Fig. 2; Supplementary Fig. 5).

**Figure 2.**
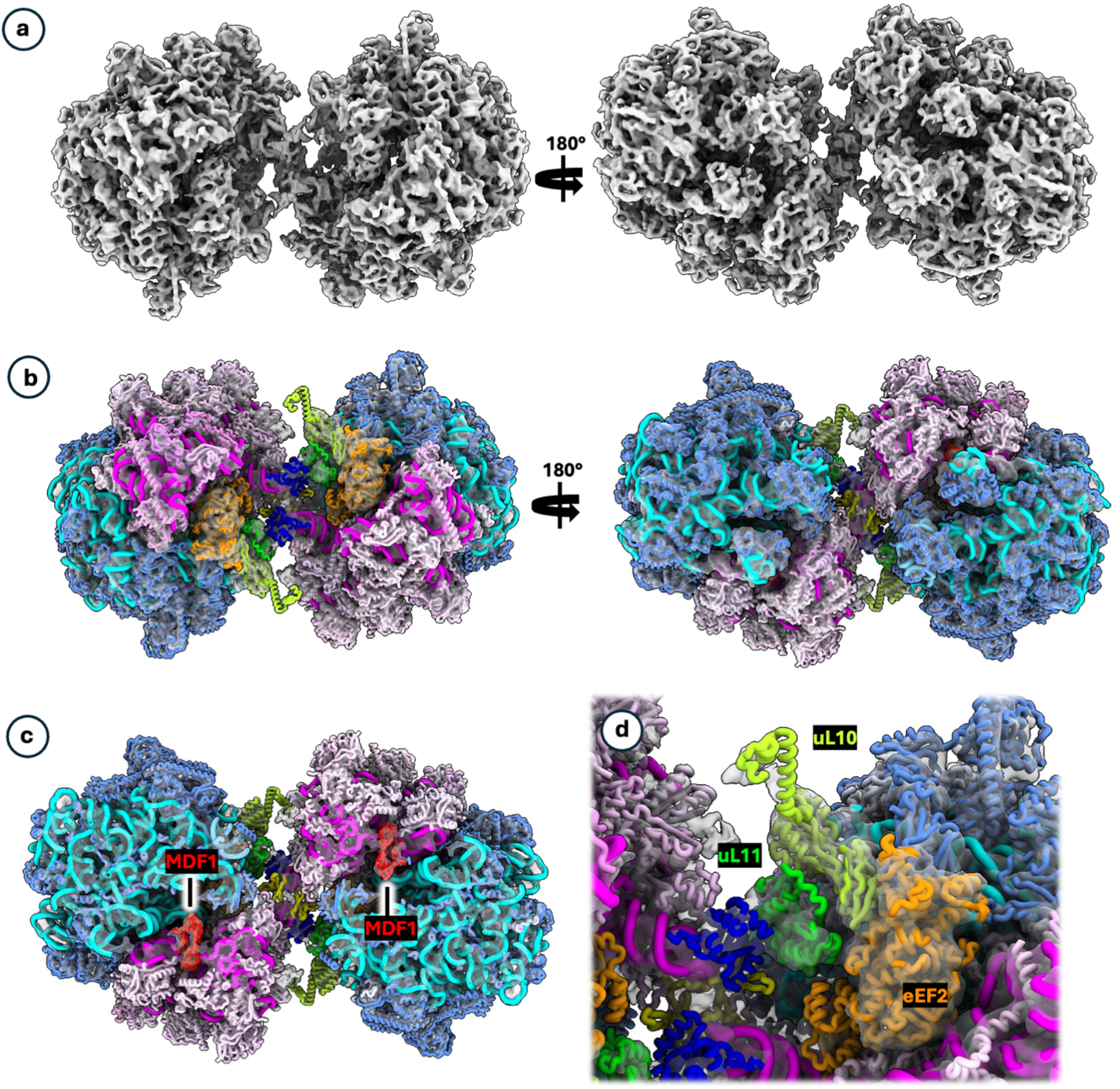
6.5 Å structure of the mHRD. **a**, dimer map (composite from two half dimers) shown in two different views, rotated by 180°. **b**, atomic model of the dimer (in liquorish representation) fitted into the map (rotated orientations as in a). Cornflower blue, large ribosomal subunit (LSU) proteins; cyan, LSU ribosomal RNA; light pink, small ribosomal subunit (SSU) proteins; magenta, SSU ribosomal RNA; shades of green, P-stalk subunits uL10 and uL11; orange, eEF2. **c**, cross-section through the E-site, showing the location of MDF1. **d**, closeup of the P-stalk and eEF2.

The refined half-dimer maps were then fitted into the dimer map, producing a high-resolution map of the mHRD (Fig. 2a). This map largely confirmed our previously published model of the mHRD from fired polar tubes (PDB-8P60; ^12^) and thus revealed that the composition and structure of the mHRD does not change significantly upon polar tube firing (Fig. 2b). Notably, both ribosomes within the dimer were bound to the hibernation factor MDF1 (Fig. 2c), stabilising the L1 stalk in a closed conformation that prevented tRNA binding at the E-site, effectively rendering the ribosomes translationally inactive.

Unlike the hibernating ribosomes of *Vairimorpha necatrix*, where an additional hibernation factor was reported to occupy the polypeptide exit tunnel (PET) ^11^, no such factor was observed in *S. lophii* mHRDs. However, several densities were now better resolved in the map of the spore-borne mHRD. Correlating prior mass spectrometry data^12^, structural models from the Protein Data Bank (PDB), and *S. lophii* homologue predictions via AlphaFold allowed us to confidently identify the corresponding proteins. We found that the eukaryotic elongation factor eEF2 is bound to the A-sites of both 70S ribosomes within the mHRD, sequestering the Sarcin/Ricin Loop (Fig. 2d; Supplementary Figure 6).

Additionally, we were able to model the ribosomal protein uL11 at the base of uL10, forming the P-stalk of both ribosomes (Fig. 2d, 3a). The inclusion of uL11 extends our knowledge of the mHRD dimer interface. Our previous work established that the primary dimer interface involved interactions between the beak regions of both ribosomes, specifically helix 52 of the SSU rRNA along with ribosomal proteins eS31 and eS12 (Fig. 3a,b ^12^). Our updated model now reveals a secondary bridge across the dimer, formed by the P-stalk bases (2×uL10 and uL11) alongside two copies of eS12 and eS31, linking the large ribosomal subunits (Fig. 3a,b). Notably, eEF2 resides at the base of the P-stalk, establishing additional contacts between uL11, LSU and SSU, thus providing additional stability to the dimer interface (Fig. 2d).

**Figure 3.**
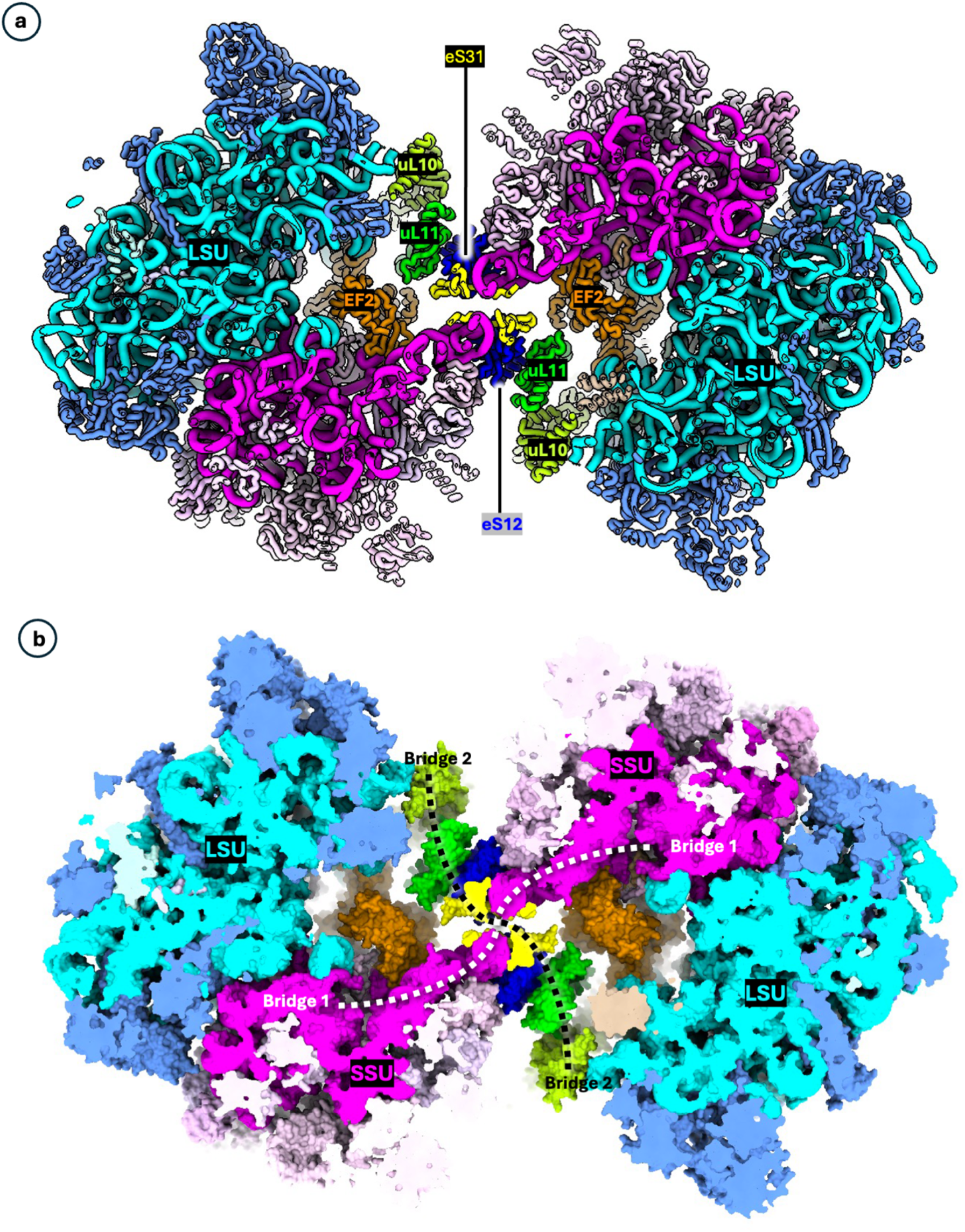
Model of the dimer interface of the mHRD. **a**, cross-section of the atomic model, showing the dimer interface subunits eS12, eS31, uL11 and Ul10 in liquorish representation. **b**, atomic model in solid representation. The mHRD is held together by two molecular bridges. one spanning the two SSUs (SSU-eS12/eS31-eS31/eS12-SSU), and a second, spanning the two LSUs (LSU-uL10-uL11-eS12/eS31-eS31/eS12-uL11-uL10-LSU).

Further inspection of the mHRD map at high contour levels revealed a third inter-dimer contact; an interaction between uL10 of one ribosome and an elongated, curved density extending from the SSU of the opposing ribosome (Supplementary Fig. 7). Although this density remains poorly resolved and its identity unknown, this observed interaction likely further stabilises the dimer and shields the P-stalks from the cellular environment. This arrangement may also explain why the P-stalks, typically flexible and thus poorly resolved in single-particle analyses, are well defined in our sub-tomogram averages.

In contrast to the HRDs, the resolution of the monosome maps obtained through our sub-tomogram averaging was insufficient to clearly visualise molecular details in their A,P, and E sites. Consequently, we were unable to resolve hibernation factors or potential tRNAs, leaving it unclear whether the monosomes represent a distinct class of hibernating ribosomes or a small pool of translationally active ones.

To further investigate the structural organisation of polar tube-associated ribosome crystals, we expanded the mask around the sub-tomogram average of the mHRD and performed 3D classification. This approach revealed distinct classes containing an array of ribosomes, in which the central density clearly corresponded to an mHRD (Supplementary Fig. 8). The results indicate that the mHRD is the fundamental unit of the crystal, similar to what has been reported by Sharma *et al* ^29^.

To model the crystal lattice, we used the auto-fitting function in ChimeraX ^31^, first superimposing our high-resolution mHRD maps and second the corresponding atomic models across all dimeric densities. This approach generated a molecular model of the ribosome crystal (Fig. 4), consisting of nine mHRDs. The derived crystal unit cell parameters were approximately 292 Å for a, 400 Å for b, and an angle of 76° for α. The crystalline array exhibited a curved conformation, with a radius of approximately 660 Å, corresponding with measured radii of spore-borne polar tubes, which range between 500 and 800 Å. Notably, the concave face of the crystal harbours the MDF1-blocked E-sites of the ribosomes (Fig. 4), while its convex face contains the occluded P-stalks.

**Figure 4.**
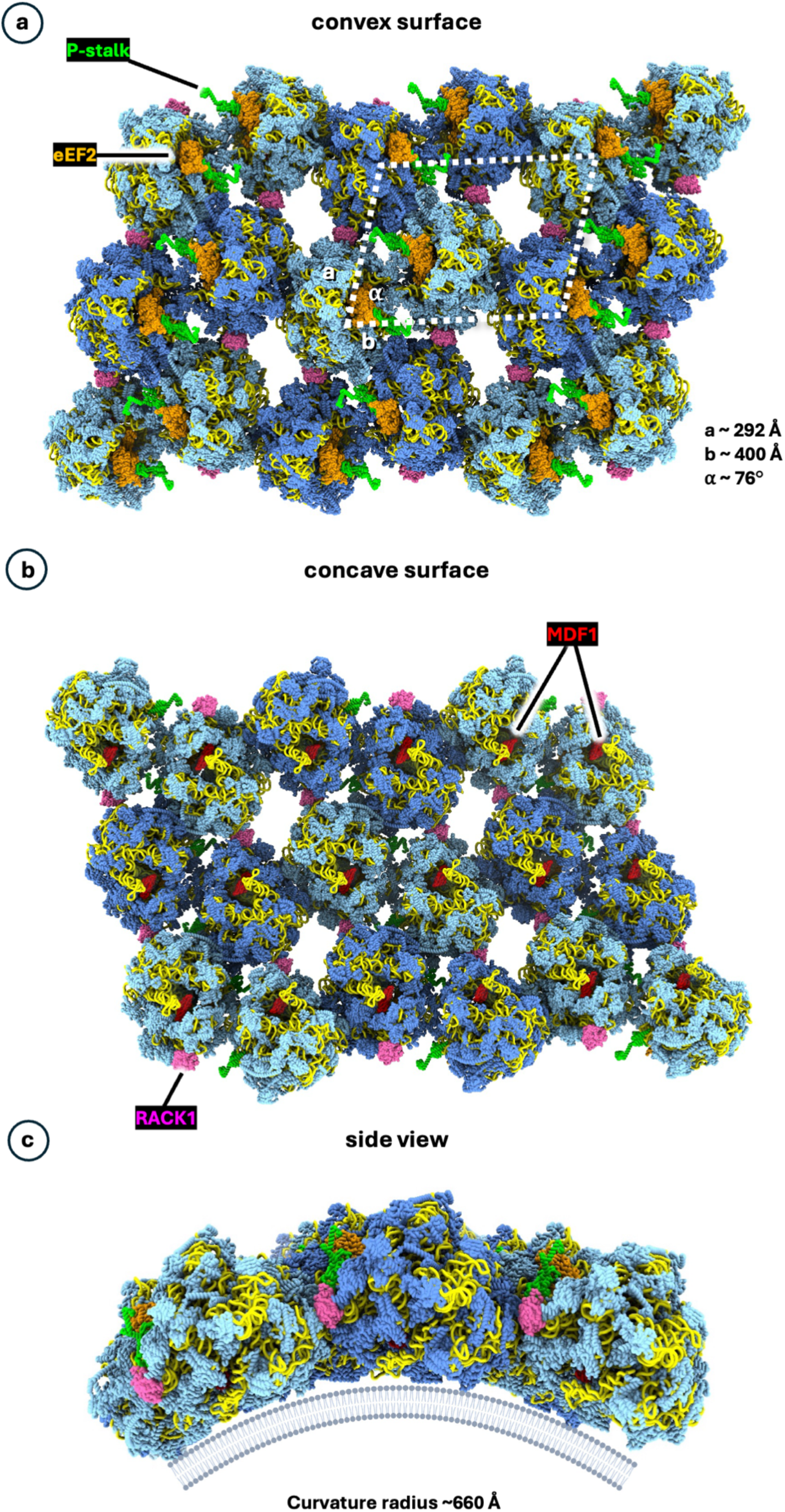
Atomic model of the polar tube-associated mHRD crystal. **a-c**, the structure of the mHRD crystal, showing its sporoplasm-facing, convex surface (**a**); the concave, polar tube-bound surface (**b**); and a cross-section (**c**). The basic unit of the crystal is the mHRD (cornflower blue / light blue). The unit cell is indicated in a. The crystal is curved, with an average curvature radius of ∼660 Å, corresponding to the polar tube’s dimensions. rRNA is shown in yellow, MDF1 in red, the P-stalks in lime, eEF2 in orange and RACK1 in hot pink.

The model revealed three distinct crystal contact interfaces between dimers. Along the short axis of the unit cell, two types of contacts were established. The first involves back-to-back interactions between large ribosomal subunits, primarily L13A, L9, L7, and L21. The second contact in this direction involves back-to-front interactions between large and small ribosomal subunits, specifically L4, L26, and Rack1. Along the long axis of the unit cell, additional back-to-back contacts between large and small ribosomal subunits were identified, including S1, S7, S27, L3, L27, L34, and ribosomal RNA. From these data it appears that the embedding of the ribosomes in the crystal does not only serve as a way to store fully assembled ribosomes but also to obstruct and immobilise additional ribosomal factors involved in ribosome biogenesis, translational fidelity and regulation, as well as various stress response mechanisms.

## Discussion

In this study, we advanced our understanding of ribosome hibernation in eukaryotes by investigating dormant *S. lophii* spores using cutting-edge cryoET. Our analysis resolved the *in-situ* structure of the mHRD at 6.5 Å, revealing the P-stalk subunits uL10 and uL11, along with eEF2, as so-far unresolved key components of the dimer interface.

The ribosomal subunits involved in the mHRD’s dimer interface are conserved across eukaryotes. We therefore hypothesise that ribosome dimerisation may be a broadly conserved mechanism among eukaryotic organisms that encounter stress conditions or undergo prolonged periods of dormancy or quiescence. Supporting this hypothesis, numerous bacterial species have been shown to form hibernating ribosome dimers under nutrient-limiting conditions ^13–21,32^. However, in bacteria, the dimer interface is established through distinct ribosome-associated factors, such as the hibernation-promoting factor (HPF) and ribosome modulation factor (RMF). Most recently, dimerization of archaeal hibernating ribosomes was also reported, where archeal hibernating ribosome dimers (aHRDs) are established by distinct factors, called aRDF-A and aRDF-B ^33^. Despite these structural differences, the presence of ribosome dimers in archaeal, bacterial and eukaryotic species suggests that dimerisation is a fundamental mechanism for ribosome hibernation, having evolved independently in these domains of life ^12^. Indicative findings from rat glioma cells ^26^ as well as mitochondria associated hibernating ribosomes in yeast ^34^, highlight that the phenomenon of ribosome oligomerisation indeed extends beyond microsporidia and is a common feature of cellular survival strategies.

Our structure of the microsporidian mHRD reveals that the P-stalk is integral to the dimer interface. The P-stalk is positioned on the large ribosomal subunit and consists of multiple proteins, including uL10 and uL11, and flexible rRNA extensions that create a dynamic platform for recruiting and stabilising translation factors ^35,36^. In active ribosomes, the P-stalk acts as part of the GTPase activating centre (GAC), facilitating the hydrolysis of GTP by translation factors, such as the eukaryotic elongation factor 2 (eEF2). The P-stalk is therefore crucial in driving key steps of translation, such as elongation and termination ^35,36^.

The fixation and occlusion of the P-stalk in the hibernating ribosome dimer suggest a mechanism to inhibit translation factor recruitment and activity. By blocking access to the P-stalk, the ribosome essentially “locks down” its GAC interface; preventing elongation factors like eEF2 from carrying out the GTP hydrolysis necessary for translocation. The P-stalk’s obstruction would reinforce the ribosomal hibernating state, ensuring that the ribosomes remain dormant until conditions allow the resumption of translation. For microsporidia, full ribosome reactivation would likely ensue when the parasite enters the host cell and starts exploiting the host’s energy metabolism to fuel its own protein biosynthesis machinery. Moreover, the arrest of the P-stalk could also serve as a protective measure to prevent premature activation of the ribosome while the microsporidian spore remains dormant. This would be crucial for maintaining the energy homeostasis in microsporidia during dormancy.

The mHRD’s dimer interface also contains eEF2, which is located at the base of the P-stalk. eEF2 is a eukaryotic elongation factor that facilitates the translocation step during protein synthesis. Once a peptide bond is formed between the amino acids on the ribosome, eEF2 uses GTP hydrolysis to move the mRNA-tRNA complex along the ribosome, shifting the ribosome into its next codon position. This action ensures that the ribosome is prepared for the next cycle of amino acid addition ^37^. eEF2 has been previously shown to be involved in the regulation of protein translation during dormancy. For example, in the cerebral cortex of hibernating chipmunks AMP-activated protein kinase (AMPK) is activated, leading to the phosphorylation and suppression of mTOR1. This cascade ultimately activates eEF2 kinase (GCN2), which phosphorylates and inactivates eEF2, suppressing protein synthesis ^38^. It has been shown that eEF2 phosphorylation can also ensue as a result of amino acid starvation via the Integrated Stress Response (ISR) in mice ^39^. As microsporidia are unable to produce many primary metabolites such as amino acids and nucleotides, they must be imported while microsporidia are inside the host cell ^40^. In their environmental spore state, however, microsporidia will run low on amino acids ^40^. Similar to the situation in mammals, this depletion could be sensed by the ISR, potentially triggering ribosome inactivation through eEF2 phosphorylation.

A study of hibernating ribosomes isolated from oocytes from Xenopus and Zebrafish showed that eEF2 is a key player in ribosome hibernation, working in conjunction with Habp4. eEF2 was found to be trapped in an inactive conformation at the ribosome’s A-site due to the binding of Habp4. Habp4 sequestered eEF2, preventing it from interacting with the tRNA-mRNA complex, thereby blocking ribosomal translocation ^41^. In our mHRD map, a Habp4-like protein is not revealed. However, the presence of eEF2 in the ribosome’s A-site suggests a comparable eEF2-mediated ribosome hibernation mechanism in microsporidia. Its emerging role in hibernating ribosomes across distantly related species suggests that the eEF2-mediated hibernation mechanism may be common in eukaryotes.

Ribosomes inside the ungerminated microsporidian spore are organised, partially as crystals, on and around the convex surface of the PT. The crystals are reminiscent of 2D arrays of ribosomes that have been reported in cold-adapted chicken embryos ^42^ or lizard oocytes ^43^. In microsporidia, the crystals adopt a curved superstructure, wherein the ribosome dimers face the PT surface via their L1 stalks, covering the E-site of the ribosome. The E-site is also the binding site for the hibernation factor MDF1, and we have previously shown that this binding leads to a closing over of the L1 stalk, rendering the ribosome inaccessible to tRNAs ^12^. Orientating the ribosome’s E-sites against the PT surface likely further decreases their accessibility to the cytosol and prevents MDF1 from dissociating.

The extensive crystal contacts in the ribosome dimer also occlude and immobilise additional ribosomal factors that are crucial for its biogenesis, fidelity, regulation and function, most notably Receptor for Activated C Kinase 1 (Rack1). This conserved protein integrates signal transduction with ribosome function. As part of the eukaryotic 40S ribosomal subunit, Rack1 is positioned near the mRNA exit channel, enabling interactions with translation factors, signalling molecules, and stress-response pathways ^44^. It regulates translation initiation, facilitates selective mRNA translation, and plays a role in ribosome biogenesis and quality control. During stress or infection, Rack1 mediates the translation of stress-specific mRNAs and can be exploited by pathogens to hijack host protein synthesis. Its multifunctional roles make it essential for linking cellular signalling and protein synthesis ^44^. Its occlusion within the crystal likely causes it to be inaccessible to any of these cytosolic factors, leading to a complete shutting down of its function.

The close association of mHRD crystals with the polar tube suggests that they are specifically tethered to its surface. However, the limited resolution of our ribosome lattice map does not reveal any distinct densities corresponding to such tethers. Future studies focusing on the ribosome arrays at higher resolution will have to be performed to provide deeper insights into how the mHRDs are anchored to the polar tube. Nevertheless, our observations indicate that crystal formation compounds ribosome hibernation in microsporidian spores and beyond. Sequestering ribosomes within a crystal would further restrict conformational changes and prevent disassembly, ensuring their structural integrity until reactivation is required.

After the PT is fired, ribosome crystals have been seen, albeit rarely, in polar tubes of the microsporidian species *V. necatrix* ^45^ and they completely disappear in our model species *S. lophii* (Fig. 1g; Supplementary Figure 3)^12^. Instead, mainly free-floating HRDs and some monosomes are observed. The ratio between mHRDs and monosomes remains unchanged (70% : 30%), indicating that the crystals mainly disassemble into free-floating mHRDs. We also find that following PT-firing, the ribosomes relocate from the outside of the PT into its lumen, supporting the model that during firing, the PT everts (like a finger of a glove)^46^. Indeed, the ribosomes exemplify how PT eversion drags the entire cytoplasmic content (sporoplasm) from the spore capsule into the PT, to ultimately catapult it into the target host cell (Fig. 5).

**Figure 5.**
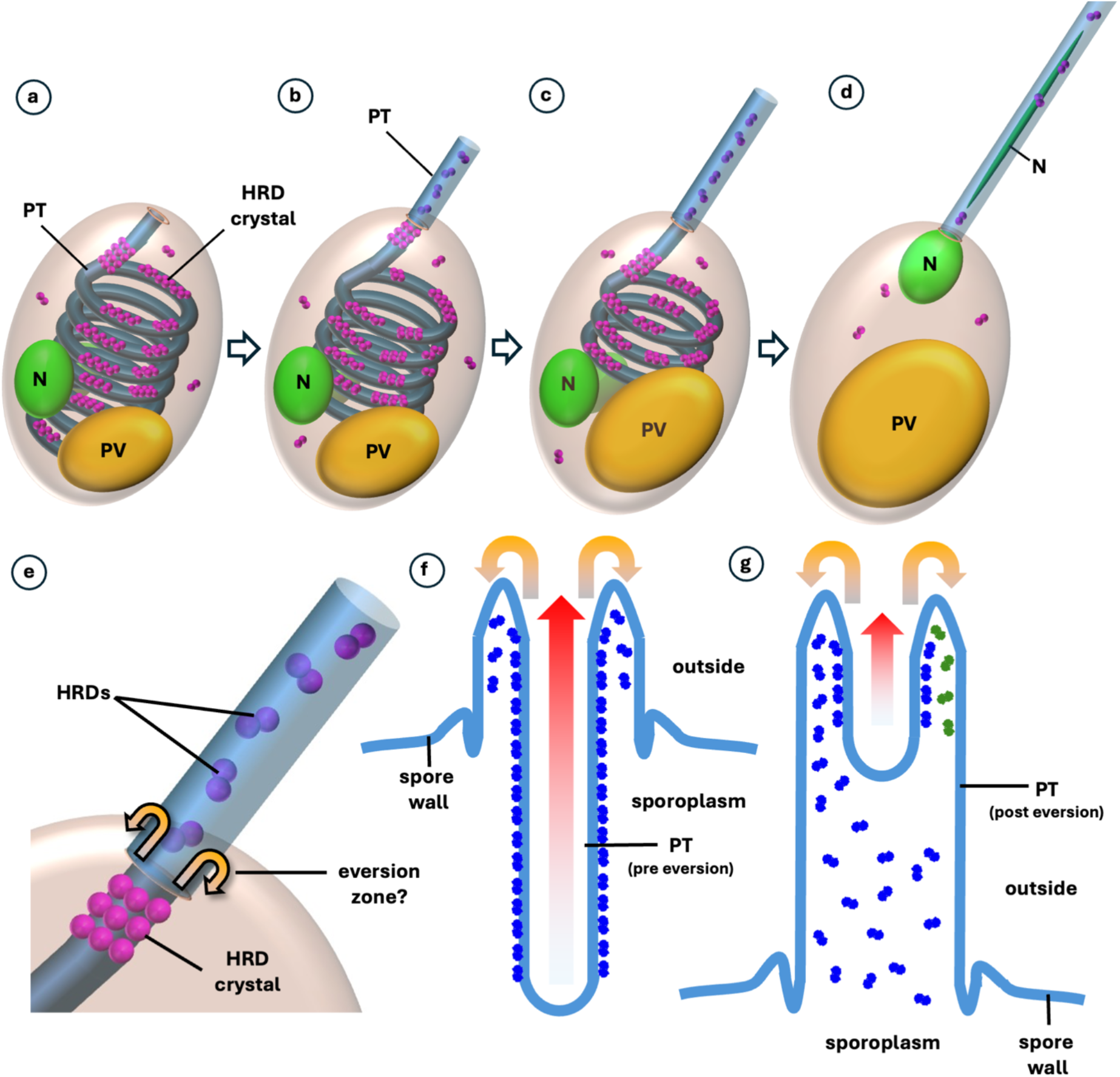
Model of ribosome reorganisation during polar tube firing. **a**, ungerminated spore. N, nuclei; PV, posterior vacuole; PT, polar tube. **b-d**, the spore germinates. The PV swells and the polar tube is pushed out of the spore. As the polar tube emerges from the spore capsule, it everts, dragging all cytoplasmic content (including ribosomes and organelles) inside and through itself. In the process, mHRD crystals, which in the spore are associated with the outer, convex surface of the polar tube, are pulled into the polar tube’s lumen. This also detaches the HRDs crystals from the PT surface and disassembles them into free-floating mHRDs. **e**, closeup of the zone where the extruding PT emerges from the spore capsule. The eversion likely takes place at the interface between capsule and outside (similar to a finger of an everting glove). **f, g**, model of the eversion, showing a cross-section of the polar tube sin two stages of germination. **f**, early stage of germination. The PT has emerged slightly from the spore, but a small portion has already everted. **g**, the polar tube has been almost fully extruded and the eversion process is nearly complete. Red arrows indicate direction of polar tube extrusion; orange arrows indicate eversion.

As the ribosome crystals dissolve, the released dimers also detach from the PT surface (Figs. 5 and 6). This would unblock the E-site, increasing the likelihood for MDF1 to dissociate and providing access for tRNAs. Releasing the ribosomes from the crystals would also make eEF2 more accessible to the cytosol and increase the likelihood for its dephosphorylation and dissociation, which would free up the A-site. However, this step likely occurs after the sporoplasm enters the host cell, as we do not observe MDF1 and eEF2 dissociation in the polar tube (Fig. 6). Abolishment of the crystal would also expose key accessory sites of the ribosome (e.g. Rack1) to the cytosol, further sensitising the ribosomes to regulatory factors, stress-response pathways and ribosome-associated quality control mechanisms. Taken together, our findings suggest that PT firing triggers a first step of hierarchical ribosome unpacking, which must ultimately culminate in the monomerisation and activation of ribosomes in the infected host cell (Fig. 6).

**Figure 6.**
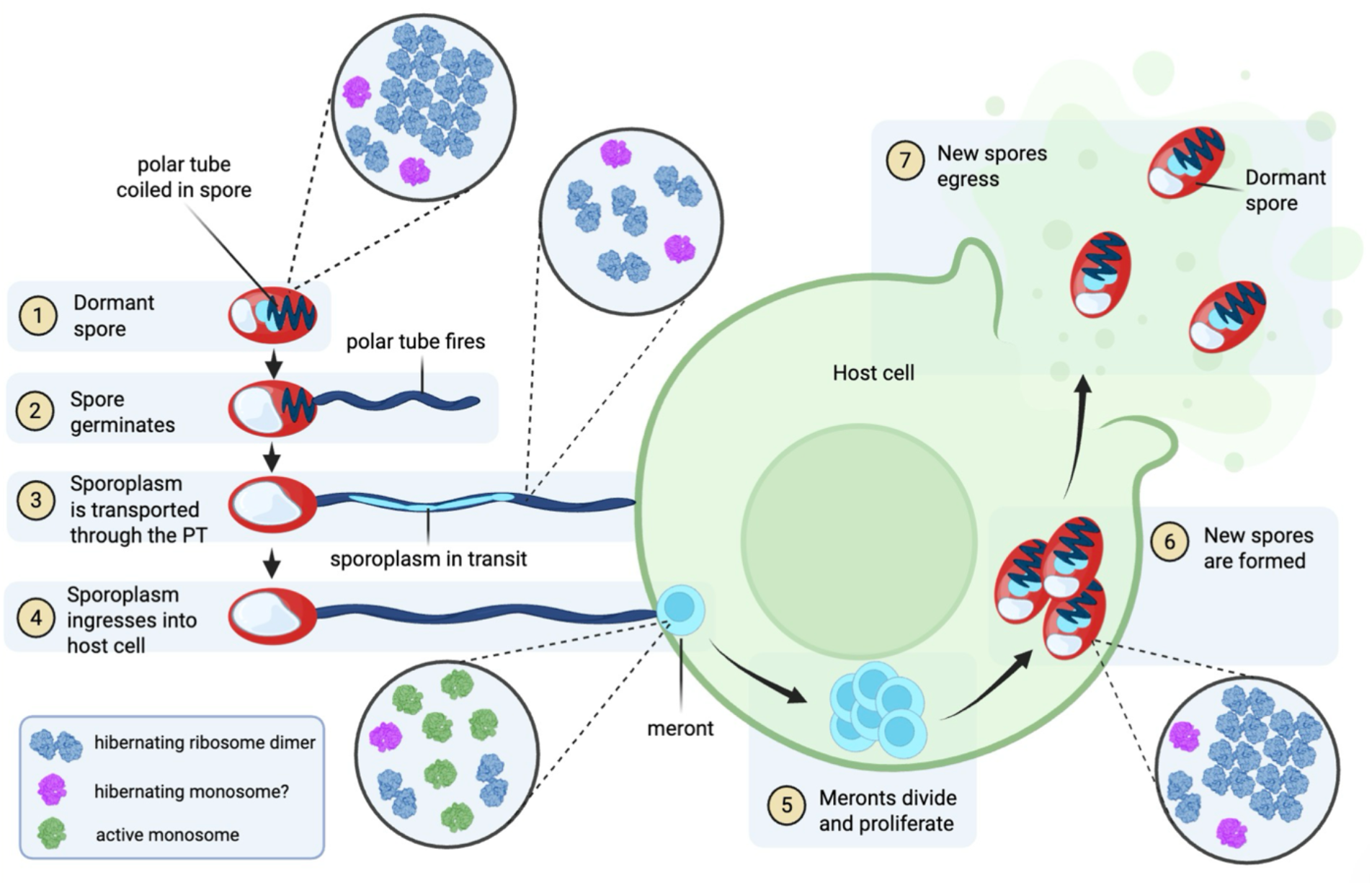
Infectious cycle of microsporidia. **(1)**, dormant microsporidian spores occur in the environment and can be ingested via contaminated food or water. Most of the ribosomes hibernate as free-floating or crystaline dimers. A subset of ribosomes exists as monosomes. These either represent a distinct hibernation state or a pool of actively translating ribosomes that maintain a baseline of translational activity. **(2)**, the spore germinates by firing the polar tube. (**3**) The polar tube harpoons a target host cell and acts as a conduit to transport the sporoplasm into the cell. The crystals have mostly disassembled into free-floating dimers. **(4)** The sporoplasm reaches the host cell and forms a meront. The ribosomes monomerise, shed their hibernation factors, and become translationally active. **(5)** The meronts divide and differentiate. **(6)** New spores are formed. The ribosomes bind to hibernation factors, dimerise, and form crystals to suppress translation. **(7)** The spores eventually egress from the host cell to disseminate further through the infected body. The figure has been created using Biorender.

## Methods

### Spore Preparation

To obtain *S. lophii* spores, clusters of cysts were harvested from monkfish (*Lophius piscatorius*) sourced from the North Atlantic and brought to shore in Brixham, Devon, UK. Microsporidia-filled xenomas were manually extracted from fish tissue. The tissue was homogenised in phosphate-buffered saline (PBS), using a scalpel until a uniform suspension was achieved. This suspension was first passed through a sterile 100 µm nylon mesh cell strainer (Fisher Scientific) to remove larger debris, and then a sterile 10 µm Pluristrainer (Fisher Scientific). Spores were purified by density gradient centrifugation using a 25-50-75-100 % Percoll gradient (Sigma) at 4°C and 3,240×*g* for 1 hour. The resulting purified spores were found in the pellet, which were then collected and subjected to three washes in sterile PBS. The final sample was stored at 4 °C with ampicillin (10 µg/ml) to prevent bacterial contamination. The concentration of spores was determined using a haemocytometer.

### CryoEM grid preparation via the Waffle Method

Purified *S. lophii* spores were pelleted by centrifugation and resuspended in as little PBS buffer as possible to form a highly concentrated suspension. 2 µl of this spore suspension were applied to the grid bar side of a glow-discharged Quantifoil R2/2 Cu 300 grid, in an assembly based on the Waffle method^27^. Prior to this, the planchette hats were treated with 1-hexadecene. The waffle assembly was high pressure frozen using a HPM Live µ machine (CryoCapCell). After freezing, the waffle assemblies were disassembled, with some falling apart immediately and others needing to be disassembled using tweezers. The grids were clipped prior to FIB milling.

### Milling of vitreous lamellae via Focussed Ion Beam Scanning Electron Microscopy (FIB-SEM)

Frozen grids were loaded for lamella milling into an Aquilos FIB-SEM (Thermo Fisher Scientific), located at the Electron BioImaging Centre (eBIC) at Diamond Light Source. An organometallic platinum layer was applied prior to milling. Using AutoTEM (Thermo Fisher Scientific), 9 areas with no ice contamination were selected for milling. Initial rough milling was performed on all target areas, followed by a polishing step. A milling angle of 20 degrees relative to the grid plane was used. The target lamella thickness was 150 nm.

### Tilt Series collection

The milled grid was loaded into a Titan Krios TEM (Thermo Fisher Scientific) with a Falcon 4i direct electron detector and a Selectris energy filter (Thermo Fisher Scientific). Using Tomo 5 (Thermo Fisher Scientific), 49 tilt series were recorded at a pixel size of 1.9 Å/px with a tilt range of –20 to +60 degrees in 2° steps, centred around the 20° milling angle. The tilt series were collected in a dose-symmetric fashion, with groupings of 3 tilts. Each tilt consisted of 8 frames, with a dose per tilt of 2.88 e^-^/Å^2^, and a total dose of 118 e^-^/A^2^ over the entire tilt series. A defocus range of –3 µm to –5 µm was used.

### Tilt Series Reconstruction and Sub-Tomogram Averaging

Motion correction and contrast transfer function (CTF) estimation were performed using Warp ^47^. Tilt series alignment was conducted with AreTomo ^48^, and tomograms were reconstructed following binning and deconvolution in Warp. Particle selection was done manually using IMOD^49^, followed by extraction in Warp with Fourier cropping at 10 Å/px resolution. A total of 25,034 particles were selected for analysis.

The EMDB entry EMD-17448 was used as an initial reference for 3D refinement. To improve the resolution of ribosome half-dimers, 3D classification and refinement were carried out in Relion^50^ until reaching the Nyquist limit. Particles were then re-extracted at 4 Å/px for further refinement, achieving a resolution of 13.6 Å. It was found that the dose weighting of tilts was incorrect and was therefore manually corrected. Particles were again re-extracted at 4 Å/px, then refined to

8.12 Å. The dataset was subsequently imported into M^47^, where image warp, volume warp and per-particle defocus refinements were performed. Initially, a temporal sampling factor of 1 was applied, followed by further refinement with a factor of 3. Following duplicate removal, an M reconstruction with no refinement was performed, followed by a per-tilt estimation of dose weight. A final reconstruction in M led to a resolution of 6.5 Å for the half-dimer structure, based on 6212 particles.

For the full dimer reconstruction, particles were centred on the middle of the dimer and re-extracted with a larger box size to capture the entire complex. This was done prior to M refinements with a temporal sampling of 3 to avoid overfitting. Poor-quality particles and monomers were removed during classification, and duplicate particles within 100 Å were eliminated. Particles were re-extracted at 3 Å/px, refined and then rotated to the symmetry axis. Using C2 symmetry in Relion^50^, the particles were refined to 7.1 Å and, with no symmetry imposed, 8.3 Å. Following 3D classification with no alignment, two strong classes were found and refined separately. The first conformation reached 9.8 Å resolution with 3109 particles, while the second conformation had a resolution of 10.9 Å with 2389 particles.

The sub-tomogram averaging workflow is illustrated in Supplementary Figure 1.

### Model building

The updated model of the spore-borne mHRD was generated by fitting our previously published structure (PDB-8P60) into the new 6.5 Å cryoEM map using ChimeraX^31^. To complete the model, structural predictions of uL10, uL11, and eEF2 were generated using Alphafold2^51^, and subsequently fitted as rigid bodies into their corresponding densities within the map. The model was then refined using Isolde ^52^. The model of the crystal was generated by first populating nine copies of the 10 Å dimer map across the sub-tomogram average of the crystal, and then fitting nine atomic models (PDB0-8P60) across these dimer maps.

## Data availability

The maps and atomic coordinates for the single particle reconstruction of the spore-borne *S. lophii* ribosome dimer were deposited to the Protein Data Bank (https://www.rcsb.org) with accession number XXXX and to the Electron Microscopy Data Bank (https://www.ebi.ac.uk/emdb) with the accession number EMD-XXXX.

## Conflict of interest

The authors declare no conflict of interest.

## Acknowledgements

BD received funding from the Leverhulme Trust (grant reference RPG-2024-417), from the European Research Council under the European Union’s Horizon 2020 research and innovation programme (grant agreement No 803894), which also funded MM, and a Wellcome Trust Seed Award (212439/Z/18/Z), which also funded BC. MM and BC were additionally funded by a Leverhulme grant (grant reverence RPG-2023-069) awarded to VG. BC received additional support from ISSF3 Wellcome Trust Institutional Strategic Support Fund (WT105618MA). We are grateful to Diamond Light Source for access and support of the cryo-EM facilities at the UK’s national Electron Bio-imaging Centre (eBIC) at Diamond Light Source, funded by the Wellcome Trust, MRC and BBRSC. We thank Ufuk Borucu at the GW4 Facility for High-Resolution Electron Cryo-Microscopy access and support. The facility was funded by the Wellcome Trust (202904/Z/16/Z and 206181/Z/17/Z) and BBSRC (BB/R000484/1). We acknowledge Lenka Koptasikova and Christian Hacker (Director of the Bioimaging Facility at the University of Exeter) for their support with High Pressure Freezing.

## Notes

### Competing Interest Statement

The authors have declared no competing interest.

## References

1. Stentiford, G. D. et al. Microsporidia – Emergent Pathogens in the Global Food Chain. Trends Parasitol 32, 336–348 (2016).

2. Han, B. & Weiss, L. M. Microsporidia: Obligate Intracellular Pathogens Within the Fungal Kingdom. Microbiol Spectr 5, (2017).

3. Han, B., Takvorian, P. M. & Weiss, L. M. Invasion of Host Cells by Microsporidia. Front Microbiol 11, (2020).

4. Wan Sajiri, W. M. H., Kua, B. C. & Borkhanuddin, M. H. Detection of Enterocytozoon hepatopenaei (EHP) (microsporidia) in several species of potential macrofauna-carriers from shrimp (Penaeus vannamei) ponds in Malaysia. J Invertebr Pathol 198, 107910 (2023).

5. Katinka, M. D. et al. Genome sequence and gene compaction of the eukaryote parasite Encephalitozoon cuniculi. Nature 414, 450–453 (2001).

6. Keeling, P. J. & Fast, N. M. Microsporidia: Biology and Evolution of Highly Reduced Intracellular Parasites. Annu Rev Microbiol 56, 93–116 (2002).

7. Texier, C., Vidau, C., Viguès, B., El Alaoui, H. & Delbac, F. Microsporidia: a model for minimal parasite–host interactions. Curr Opin Microbiol 13, 443–449 (2010).

8. Heinz, E. et al. Plasma Membrane-Located Purine Nucleotide Transport Proteins Are Key Components for Host Exploitation by Microsporidian Intracellular Parasites. PLoS Pathog 10, e1004547 (2014).

9. Fayet, M. et al. New insights into Microsporidia polar tube function and invasion mechanism. Journal of Eukaryotic Microbiology 71, (2024).

10. Chen, Y. et al. The microsporidian polar tube: origin, structure, composition, function, and application. Parasit Vectors 16, 305 (2023).

11. Barandun, J., Hunziker, M., Vossbrinck, C. R. & Klinge, S. Evolutionary compaction and adaptation visualized by the structure of the dormant microsporidian ribosome. Nat Microbiol 4, 1798–1804 (2019).

12. McLaren, M. et al. CryoEM reveals that ribosomes in microsporidian spores are locked in a dimeric hibernating state. Nat Microbiol 8, 1834–1845 (2023).

13. Kato, T. et al. Structure of the 100S Ribosome in the Hibernation Stage Revealed by Electron Cryomicroscopy. Structure 18, 719–724 (2010).

14. Beckert, B. et al. Structure of the Bacillus subtilis hibernating 100S ribosome reveals the basis for 70S dimerization. EMBO J 36, 2061–2072 (2017).

15. Khusainov, I. et al. Structures and dynamics of hibernating ribosomes from Staphylococcus aureus mediated by intermolecular interactions of <scp>HPF</scp>. EMBO J 36, 2073–2087 (2017).

16. Matzov, D. et al. The cryo-EM structure of hibernating 100S ribosome dimer from pathogenic Staphylococcus aureus. Nat Commun 8, 723 (2017).

17. Franken, L. E. et al. A general mechanism of ribosome dimerization revealed by single-particle cryo-electron microscopy. Nat Commun 8, 722 (2017).

18. Beckert, B. et al. Structure of a hibernating 100S ribosome reveals an inactive conformation of the ribosomal protein S1. Nat Microbiol 3, 1115–1121 (2018).

19. Flygaard, R. K., Boegholm, N., Yusupov, M. & Jenner, L. B. Cryo-EM structure of the hibernating Thermus thermophilus 100S ribosome reveals a protein-mediated dimerization mechanism. Nat Commun 9, 4179 (2018).

20. Usachev, K. S. et al. Dimerization of long hibernation promoting factor from Staphylococcus aureus: Structural analysis and biochemical characterization. J Struct Biol 209, 107408 (2020).

21. Feaga, H. A., Kopylov, M., Kim, J. K., Jovanovic, M. & Dworkin, J. Ribosome Dimerization Protects the Small Subunit. J Bacteriol 202, (2020).

22. Nicholson, D. et al. Adaptation to genome decay in the structure of the smallest eukaryotic ribosome. Nat Commun 13, 591 (2022).

23. Ehrenbolger, K. et al. Differences in structure and hibernation mechanism highlight diversification of the microsporidian ribosome. PLoS Biol 18, e3000958 (2020).

24. Wells, J. N. et al. Structure and function of yeast Lso2 and human CCDC124 bound to hibernating ribosomes. PLoS Biol 18, e3000780 (2020).

25. Brown, A., Baird, M. R., Yip, M. C., Murray, J. & Shao, S. Structures of translationally inactive mammalian ribosomes. Elife 7, (2018).

26. Krokowski, D. et al. Characterization of hibernating ribosomes in mammalian cells. Cell Cycle 10, 2691–2702 (2011).

27. Kelley, K. et al. Waffle Method: A general and flexible approach for improving throughput in FIB-milling. Nat Commun 13, 1857 (2022).

28. Klykov, O. et al. In situ cryo-FIB/SEM Specimen Preparation Using the Waffle Method. Bio Protoc 12, (2022).

29. Sharma, H., Jespersen, N., Ehrenbolger, K.Carlson, L.-A. & Barandun, J. Ultrastructural insights into the microsporidian infection apparatus reveal the kinetics and morphological transitions of polar tube and cargo during host cell invasion. PLoS Biol 22, e3002533 (2024).

30. Tegunov, D., Xue, L., Dienemann, C., Cramer, P. & Mahamid, J. Multi-particle cryo-EM refinement with M visualizes ribosome-antibiotic complex at 3.5 Å in cells. Nat Methods 18, 186–193 (2021).

31. Pettersen, E. F. et al. <scp>UCSF ChimeraX</scp> : Structure visualization for researchers, educators, and developers. Protein Science 30, 70–82 (2021).

32. Wang, T. et al. Ribosome Hibernation as a Stress Response of Bacteria. Protein Pept Lett 27, 1082–1091 (2020).

33. Hassan, A. H. et al. Novel archaeal ribosome dimerization factor facilitating unique 30S–30S dimerization. Nucleic Acids Res 53, (2025).

34. Gemin, O. et al. Ribosomes hibernate on mitochondria during cellular stress. Nat Commun 15, 8666 (2024).

35. Filipek, K. et al. Phosphorylation of P-stalk proteins defines the ribosomal state for interaction with auxiliary protein factors. EMBO Rep 25, 5478–5506 (2024).

36. Liljas, A. & Sanyal, S. The enigmatic ribosomal stalk. Q Rev Biophys 51, e12 (2018).

37. Kaul, G., Pattan, G. & Rafeequi, T. Eukaryotic elongation factor-2 (eEF2): its regulation and peptide chain elongation. Cell Biochem Funct 29, 227–234 (2011).

38. Yamada, S., Kamata, T., Nawa, H., Sekijima, T. & Takei, N. AMPK activation, eEF2 inactivation, and reduced protein synthesis in the cerebral cortex of hibernating chipmunks. Sci Rep 9, 11904 (2019).

39. Harding, H. P. et al. The ribosomal P-stalk couples amino acid starvation to GCN2 activation in mammalian cells. Elife 8, (2019).

40. Dean, P., Hirt, R. P. & Embley, T. M. Microsporidia: Why Make Nucleotides if You Can Steal Them? PLoS Pathog 12, e1005870 (2016).

41. Leesch, F. et al. A molecular network of conserved factors keeps ribosomes dormant in the egg. Nature 613, 712–720 (2023).

42. Morimoto, T., Blobel, G. & Sabatini, D. D. RIBOSOME CRYSTALLIZATION IN CHICKEN EMBRYOS. J Cell Biol 52, 338–354 (1972).

43. Unwin, P. N. T. Three-dimensional model of membrane-bound ribosomes obtained by electron microscopy. Nature 269, 118–122 (1977).

44. Adams, D. R., Ron, D. & Kiely, P. A. RACK1, A multifaceted scaffolding protein: Structure and function. Cell Communication and Signaling 9, 22 (2011).

45. Sharma, H., Jespersen, N., Ehrenbolger, K.Carlson, L.-A. & Barandun, J. Ultrastructural insights into the microsporidian infection apparatus reveal the kinetics and morphological transitions of polar tube and cargo during host cell invasion. PLoS Biol 22, e3002533 (2024).

46. Chang, R. et al. Energetics of the microsporidian polar tube invasion machinery. Elife 12, (2024).

47. Tegunov, D., Xue, L., Dienemann, C., Cramer, P. & Mahamid, J. Multi-particle cryo-EM refinement with M visualizes ribosome-antibiotic complex at 3.5 Å in cells. Nat Methods 18, 186–193 (2021).

48. Zheng, S. et al. AreTomo: An integrated software package for automated marker-free, motion-corrected cryo-electron tomographic alignment and reconstruction. J Struct Biol X 6, 100068 (2022).

49. Kremer, J. R., Mastronarde, D. N. & McIntosh, J. R. Computer Visualization of Three-Dimensional Image Data Using IMOD. J Struct Biol 116, 71–76 (1996).

50. Scheres, S. H. W. RELION: Implementation of a Bayesian approach to cryo-EM structure determination. J Struct Biol 180, 519–530 (2012).

51. Jumper, J. et al. Highly accurate protein structure prediction with AlphaFold. Nature 596, 583–589 (2021).

52. Croll, T. I. ISOLDE : a physically realistic environment for model building into low-resolution electron-density maps. Acta Crystallogr D Struct Biol 74, 519–530 (2018).

