## Supplementary Figures for "New clues about ribosome hibernation in microsporidia revealed by cryo-electron tomography"

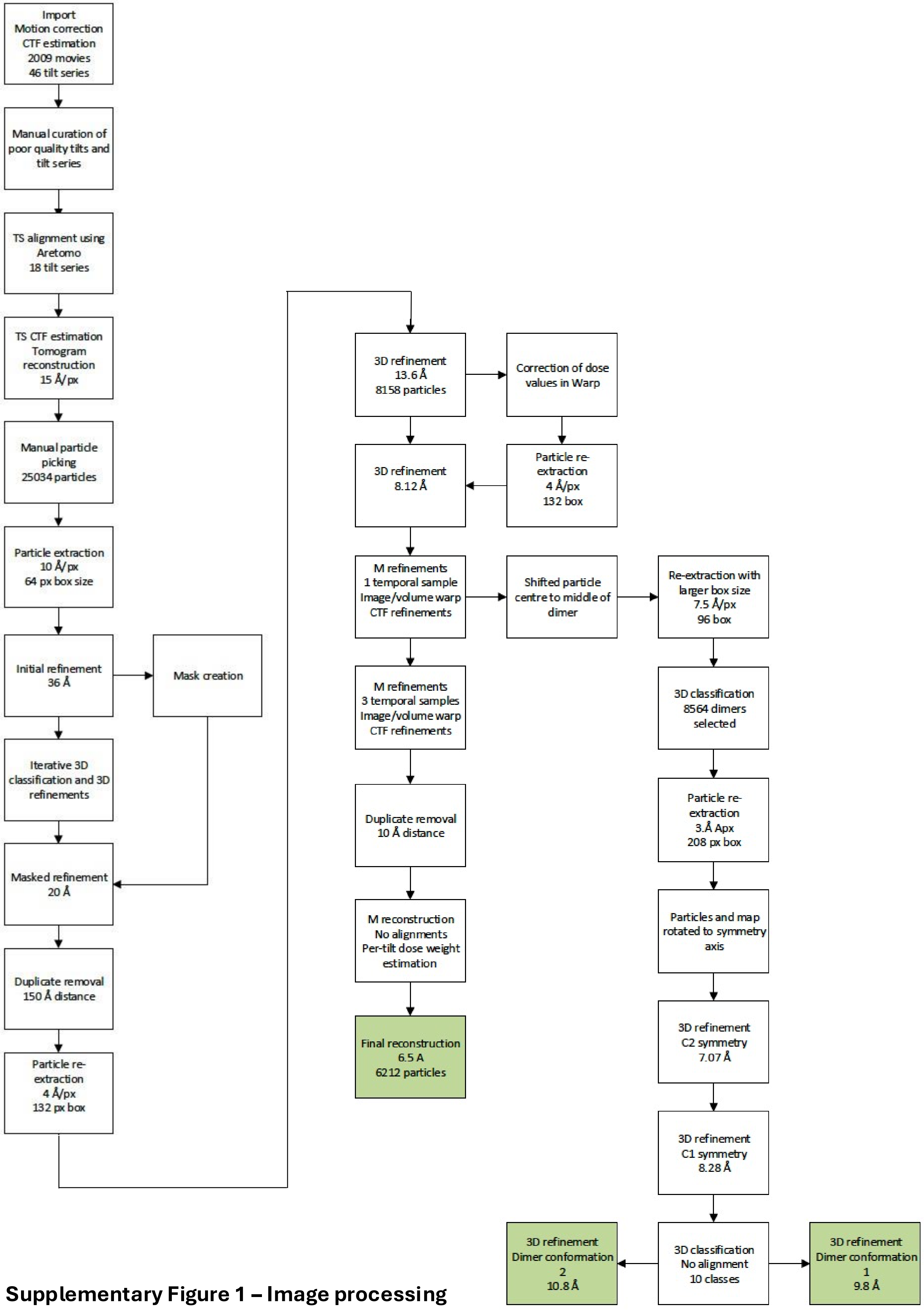


**Supplementary Figure 1 – Image processing flowchart**

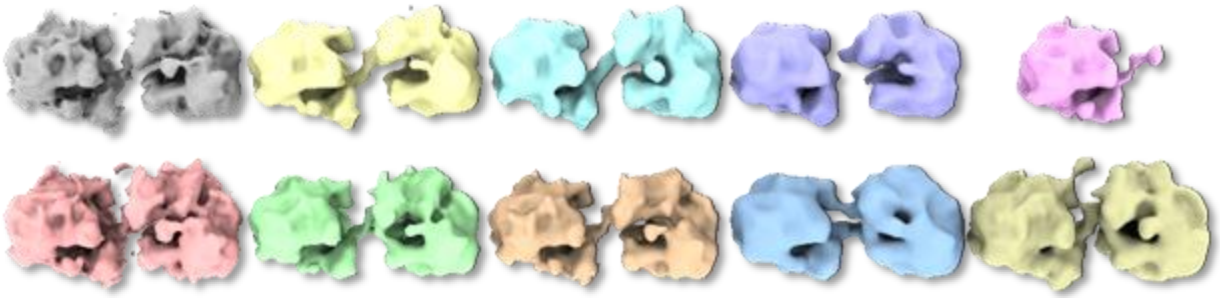

**Supplementary Figure 2 – 3D classification of ribosomes in dormant microsporidian spores**

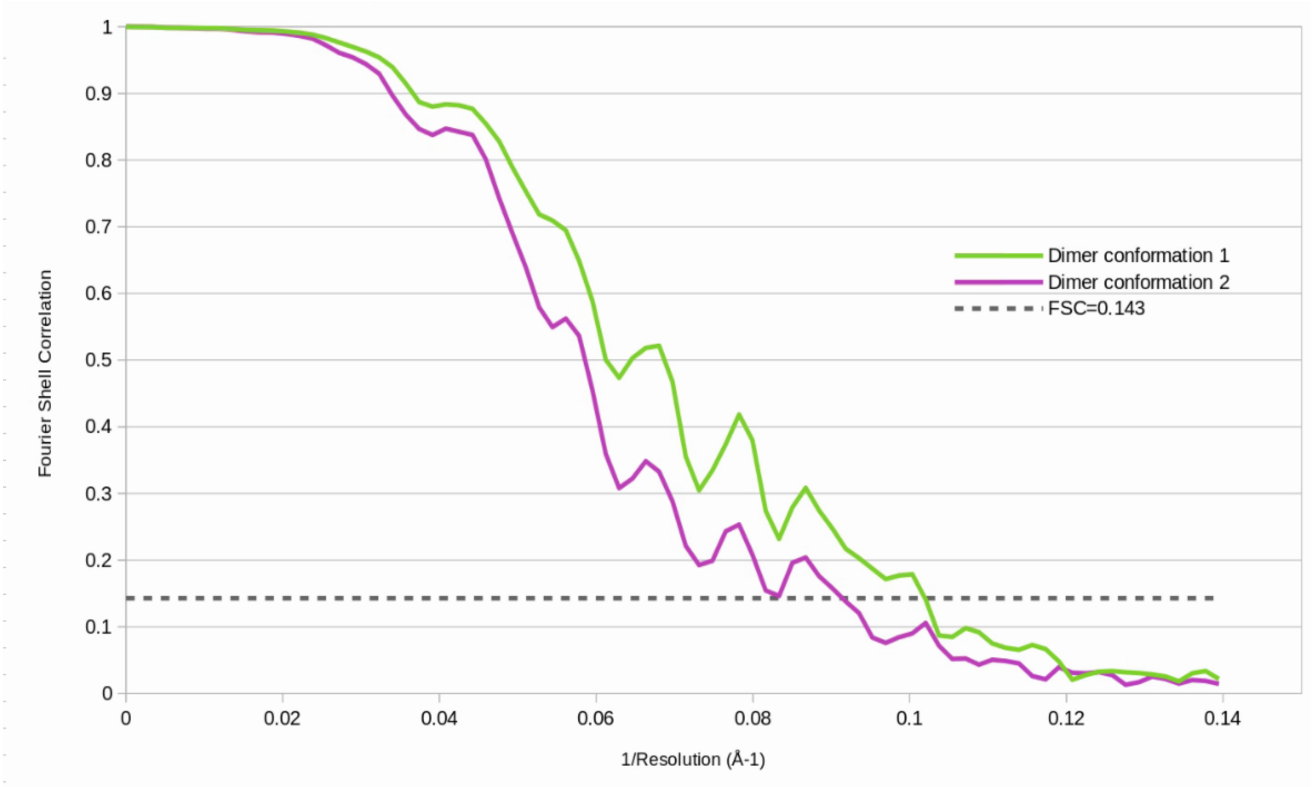


**Supplementary Figure 3 – Resolution estimation for the mHRD.**

Gold-standard Fourier Shell Correlation (FSC), indicating a global resolution of 9.8 Å for Conformation 1 and 10.9 Å for Conformation 2.

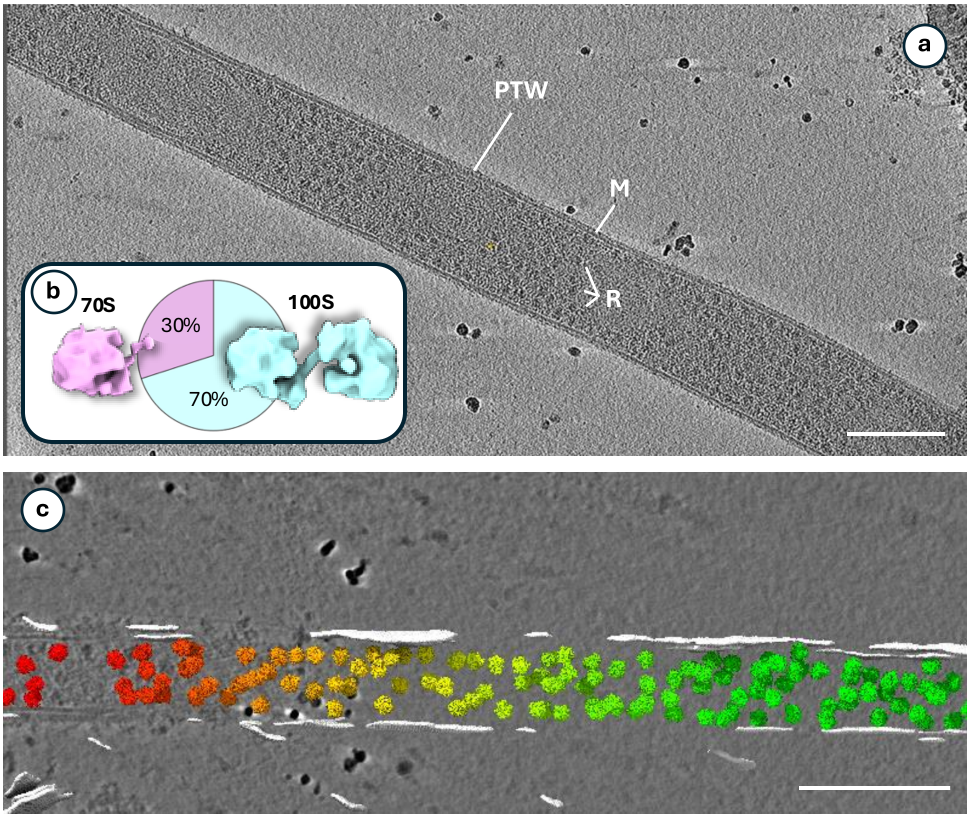


**Supplementary Figure 4 – Ribosomes in fired polar tube. a**, tomographic slice of a fired polar tube, showing multiple free-floating mHRDs (R) in the polar tube lumen. PTW, polar tube wall; M, membrane. **b**, 3D classification reveals a 70% / 30% distribution between mHRDs and monosomes. **c**, segmented tonogram of a fired polar tube showing the positions of mHDRs within it in 3D, with tomographic slice shown in the background. Scale bars, 200 nm.

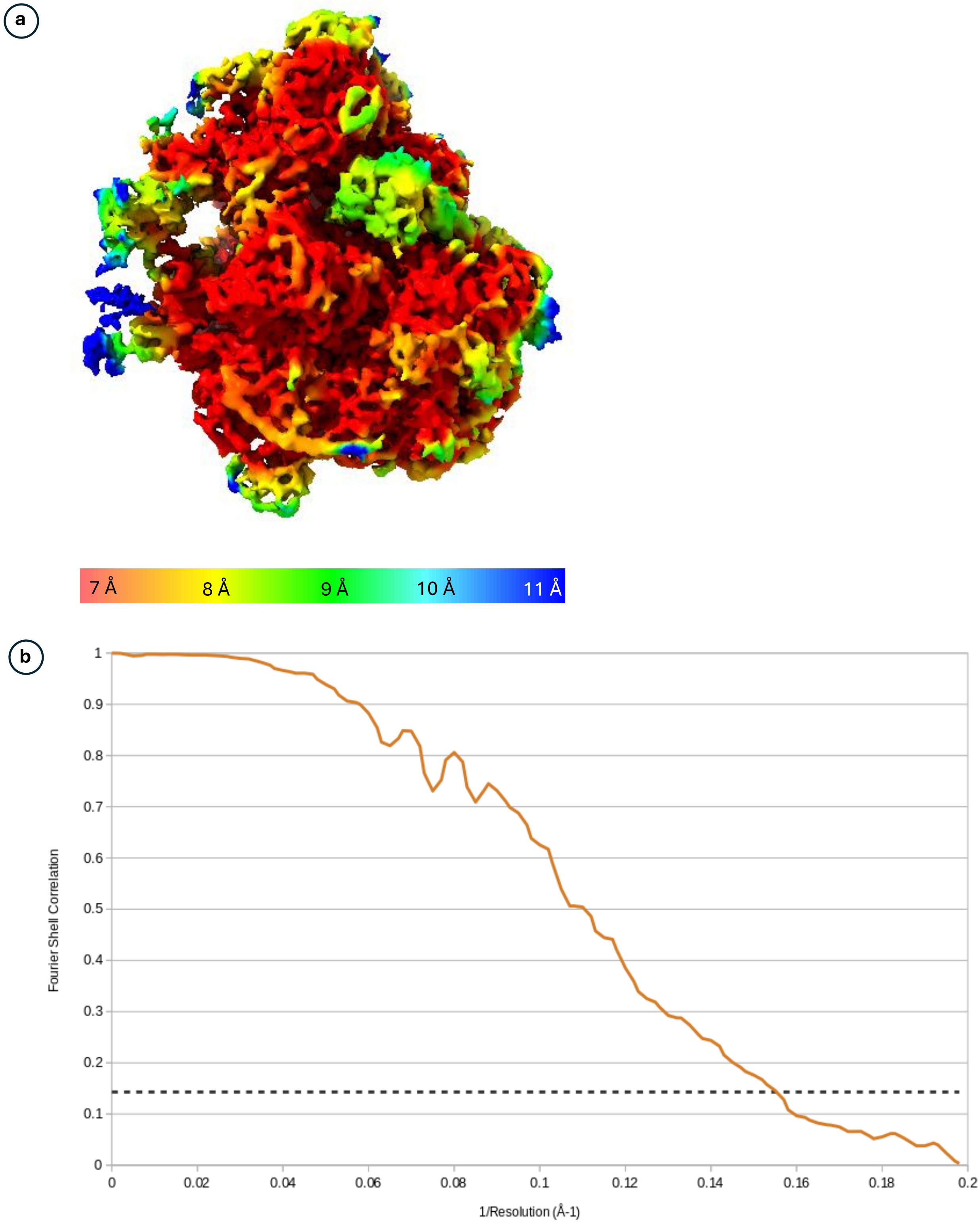


**Supplementary Figure 5 – Resolution estimation for the half-dimer.**
**a,** local resolution map; **b**, gold-standard Fourier Shell Correlation (FSC), indicating a global resolution of 6.5 Å.

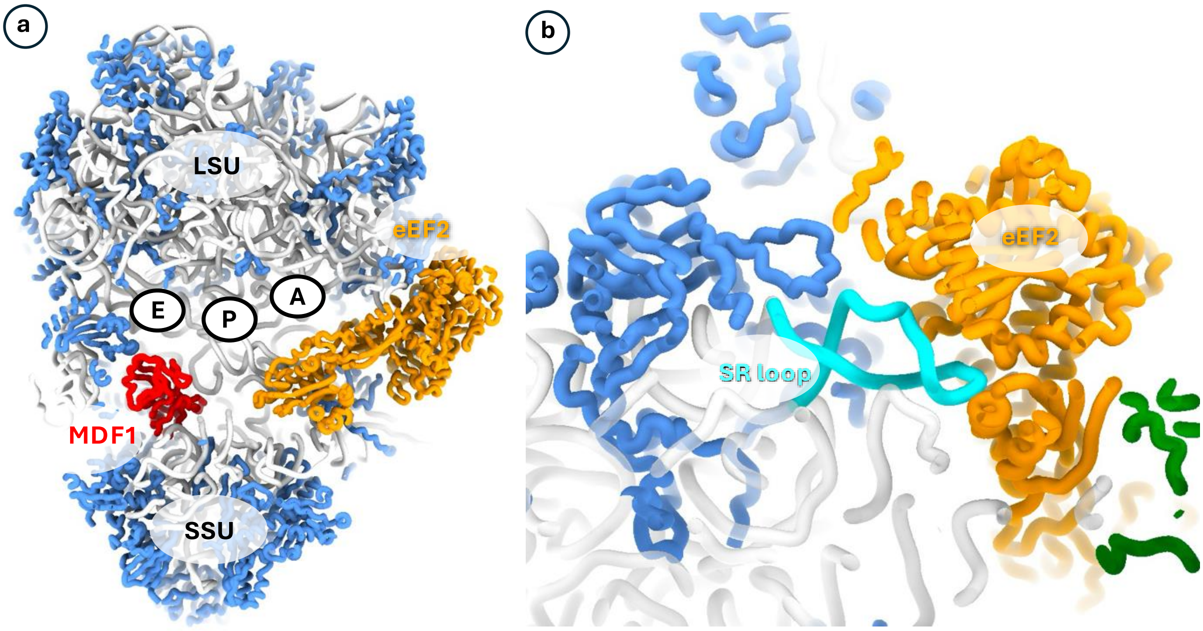


**Supplementary Figure 6 – Hibernation factors occupying A,P and E-sites**
**a**, Cross section of the mHRD, showing how MDF1 and eEF2 occupy E and A sites of the ribosome, respectively. **b**, eEF2 binding the Sarcin-Ricin (SR; light blue) loop of the large ribosomal RNA.

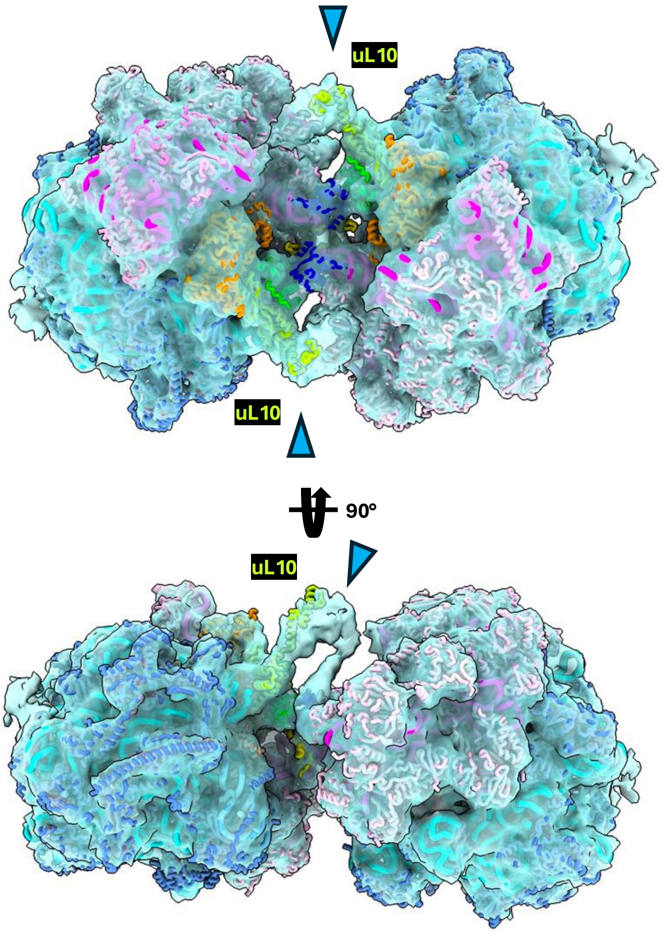


**Supplementary Figure 7 – The P-stalks form a third dimer interface**
Two views of the dimer map rotated 90 ° relative to each other. The P-stalk subunit uL10 connects with the opposite ribosome via an unidentified density.

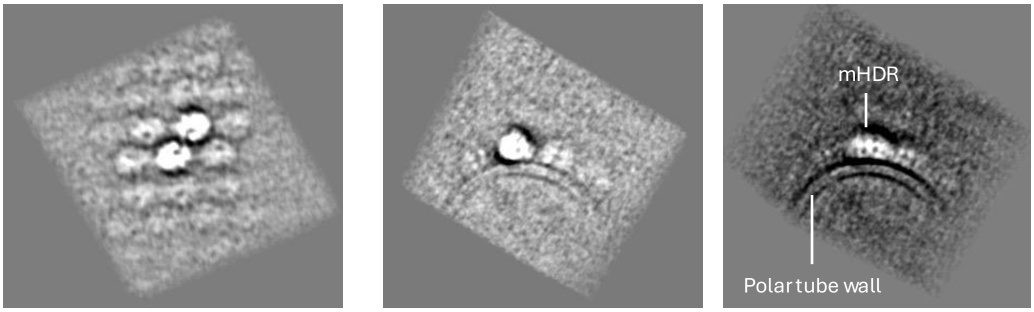


Top view
25 slices averaged

Side view
25 slices averaged

Side view
100 slices averaged

**Supplementary Figure 8 – Tomographic slices through sub-tomogram average of a polar tube-bound mHRD crystal.**
